## supplementary figures for "Maturation and detoxification of synphilin-1 inclusion bodies regulated by sphingolipids"

### Figure 2-figure supplement 1

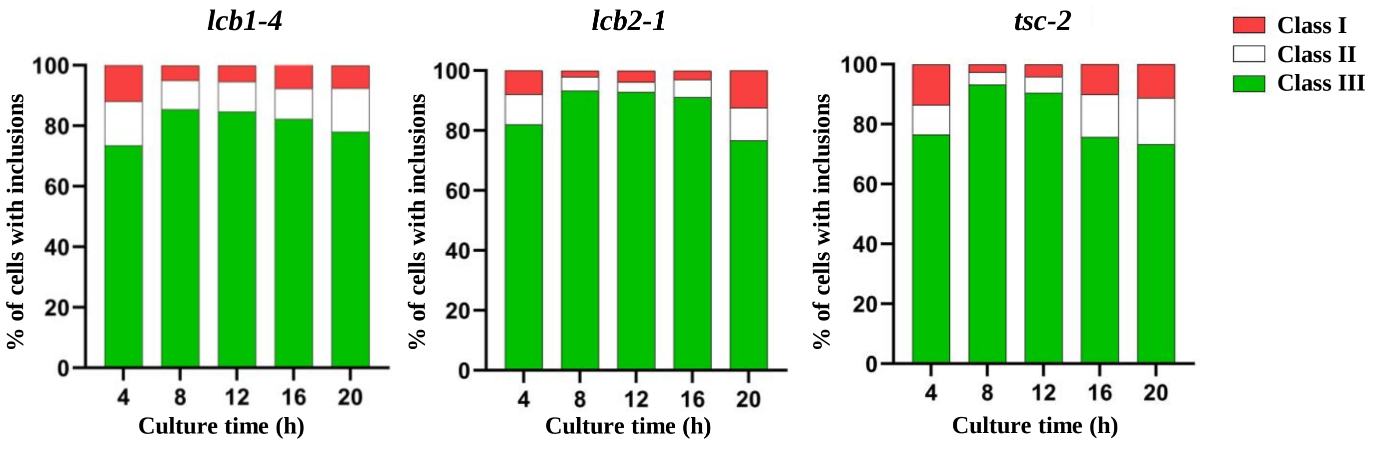

### Figure 4-figure supplement 1

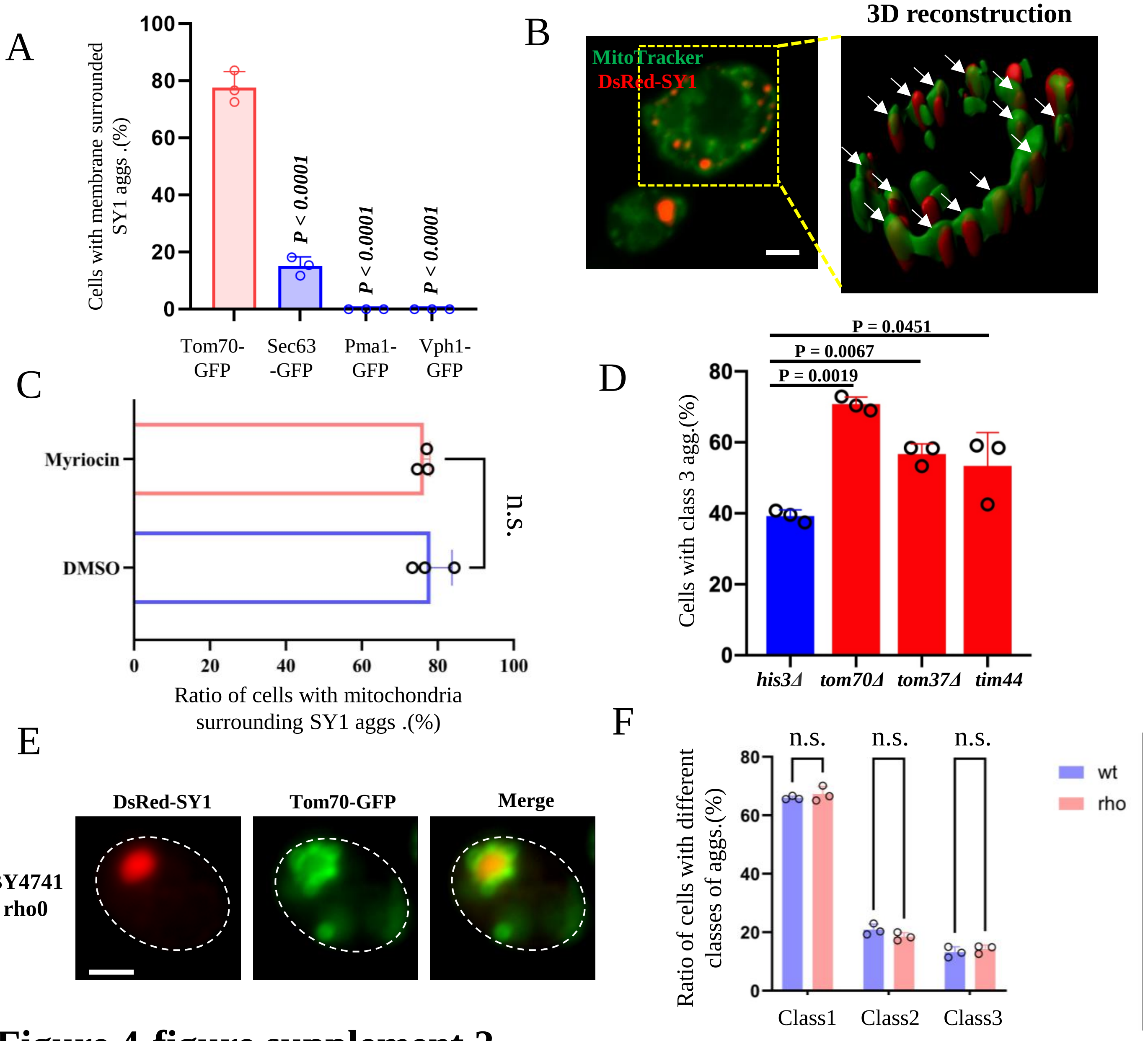


### Figure 4-figure supplement 2

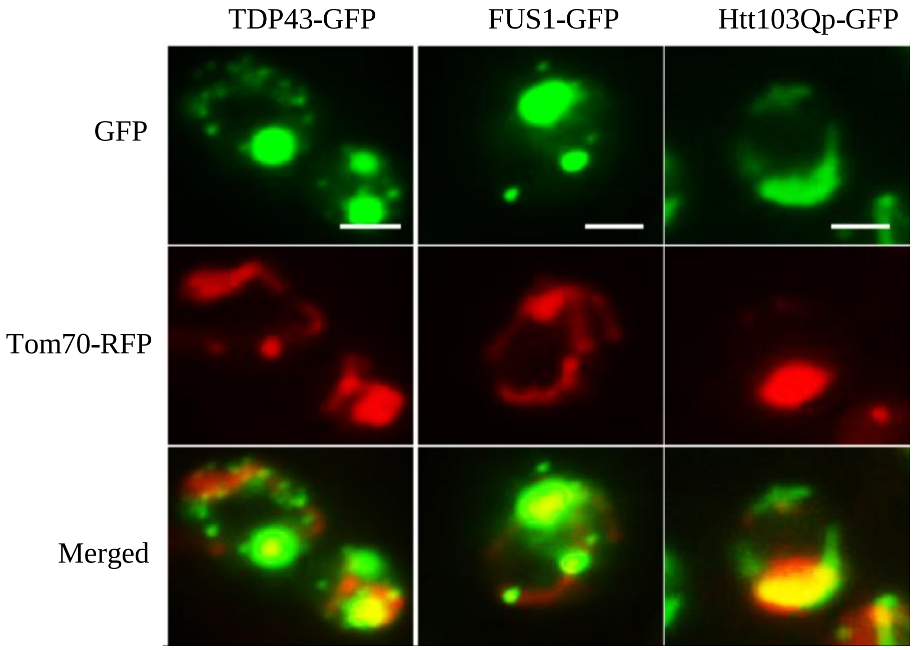

### Figure 5-figure supplement 1

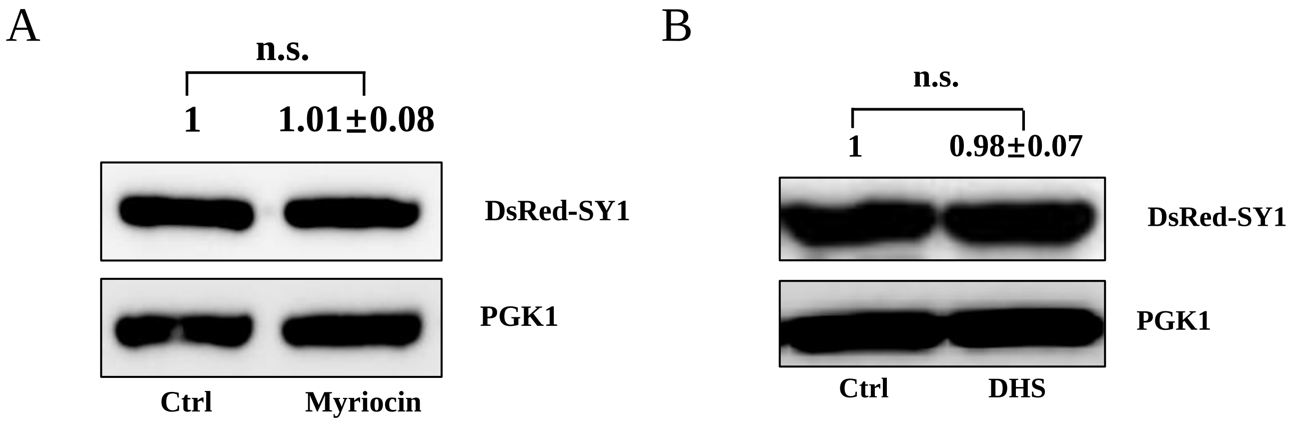


### Figure 5-figure supplement 2

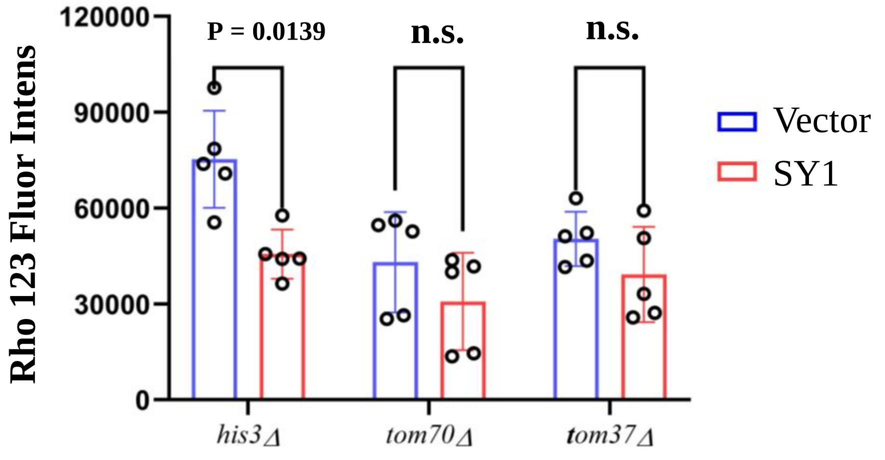

### Figure 6-figure supplement 1

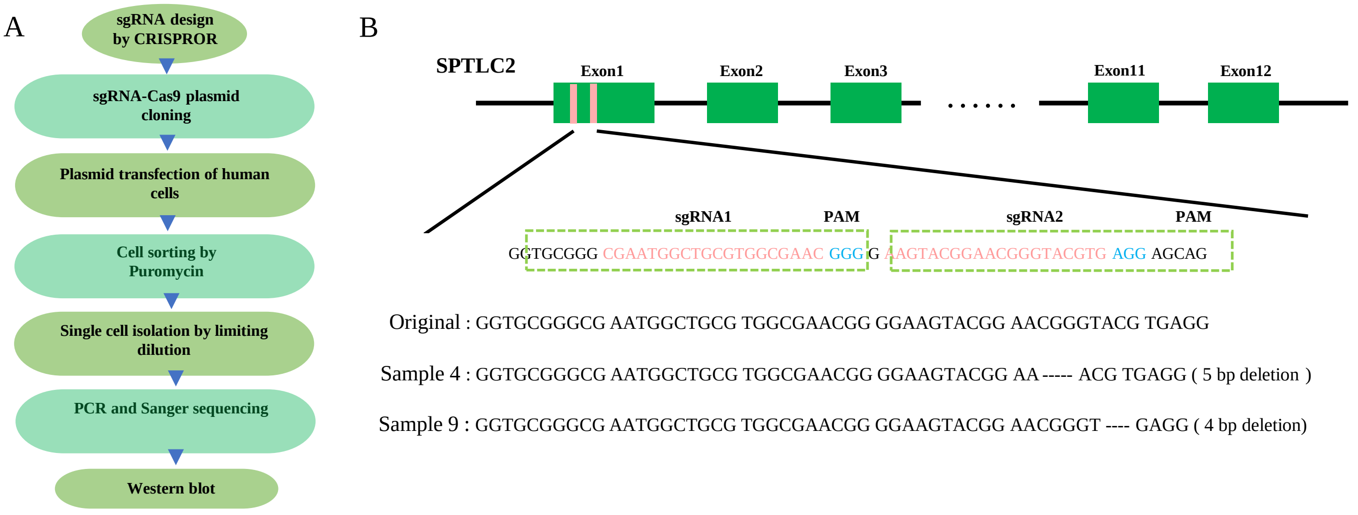


### Figure 6-figure supplement 2

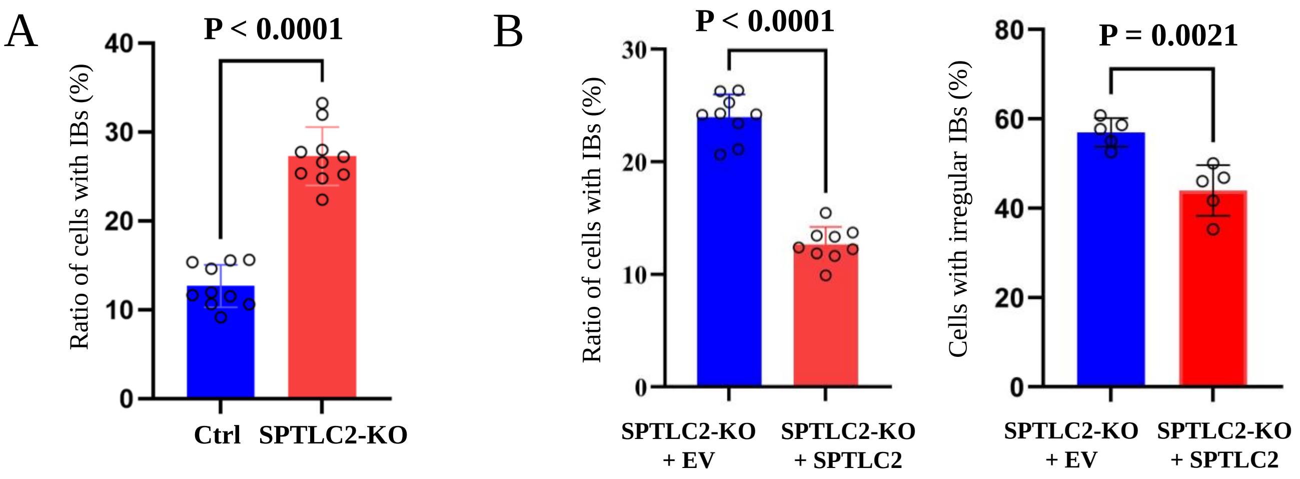


### Figure 6-figure supplement 3

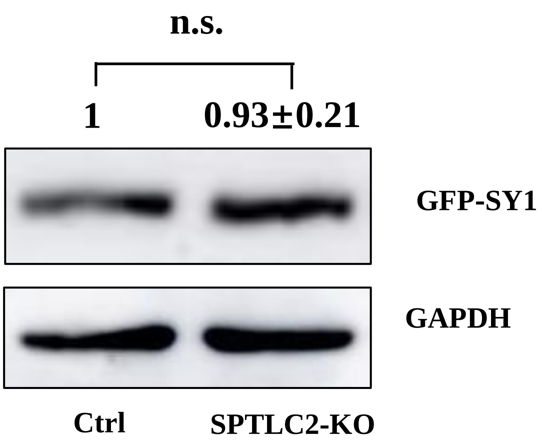

### Figure 7-figure supplement 1

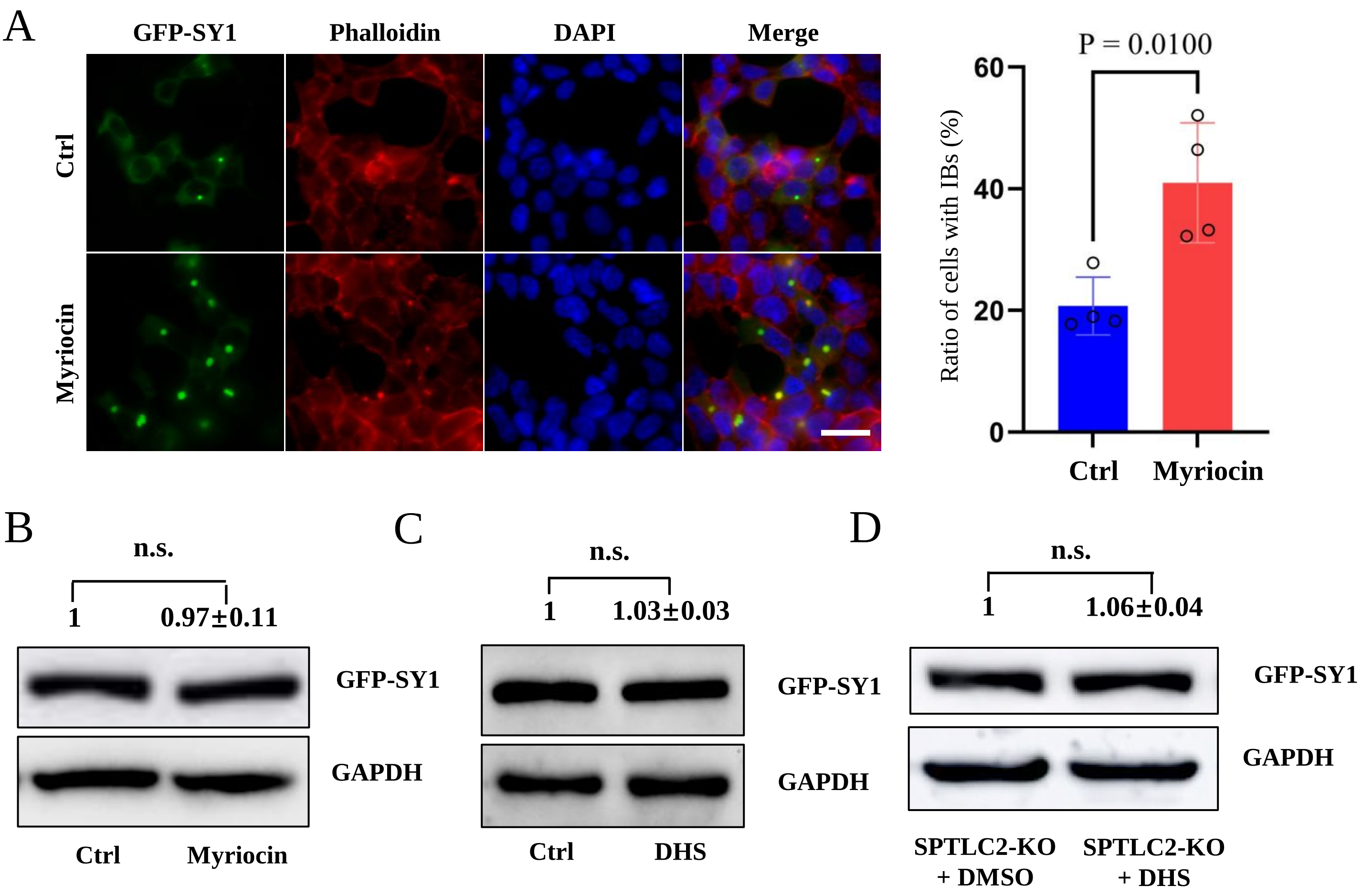


### Figure 7-figure supplement 2

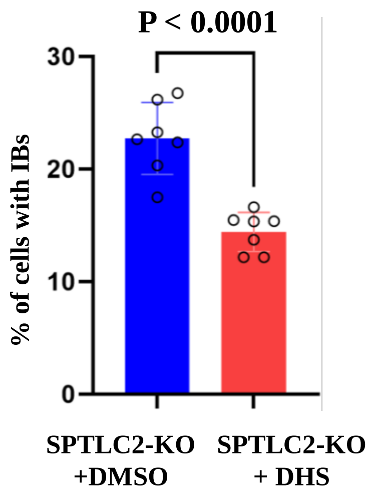
