## Supplementary file 1 for "Maturation and detoxification of synphilin-1 inclusion bodies regulated by sphingolipids"

**Table S1** Mutants with manually confirmed increased class 3 synphilin-1 aggregates

| No. | ORF | Gene | Manual-Dif % of cell with SY1 aggs in class 3 (compare to wt) | P-value |
| --- | --- | --- | --- | --- |
| 1 | YNL006W | lst8-6 | 85.8882 | 0.0011 |
| 2 | YGL007W | BRP1 | 70.6323 | 0.0001 |
| 3 | YDL126C | cdc48-4601 | 68.9313 | 0.0013 |
| 4 | YPR044C | OPI11 | 63.6510 | 0.0001 |
| 5 | YDL126C | cdc48-2 | 59.0261 | 0.0008 |
| 6 | YDL028C | mps1-1 | 58.8152 | 0.0051 |
| 7 | YER093C | tsc11-7 | 57.7372 | 0.0016 |
| 8 | YER093C | tsc11-5 | 57.4031 | 0.0052 |
| 9 | YNR010W | CSE2 | 56.1509 | 0.0045 |
| 10 | YBR058C-A | tsc3-2 | 55.5107 | 0.0044 |
| 11 | YOL081W | IRA2 | 53.2526 | 0.0004 |
| 12 | YKL203C | tor2-29 | 53.2010 | 0.0138 |
| 13 | YOL052C | SPE2 | 52.9591 | 0.0368 |
| 14 | YOL108C | INO4 | 51.9124 | 0.0128 |
| 15 | YDL213C | NOP6 | 51.4097 | 0.0023 |
| 16 | YPR024W | YME1 | 50.9979 | 0.0056 |
| 17 | YDR292C | srp101-47 | 49.1986 | 0.0045 |
| 18 | YCL045C | EMC1 | 48.9180 | 0.0436 |
| 19 | YMR235C | rna1-1 | 48.1705 | 0.0089 |
| 20 | YHR111W | UBA4 | 47.5794 | 0.0007 |
| 21 | YGR027C | RPS25A | 46.8475 | 0.0026 |
| 22 | YDR062W | lcb2-1 | 46.1388 | 0.0076 |
| 23 | YGL231C | EMC4 | 45.9566 | 0.0003 |
| 24 | YOR141C | ARP8 | 44.6984 | 0.0037 |
| 25 | YNL229C | URE2 | 44.5722 | 0.0002 |
| 26 | YGR105W | VMA21 | 44.0698 | 0.0004 |
| 27 | YHR034C | PIH1 | 43.8134 | 0.0002 |
| 28 | YJL194W | cdc6-1 | 43.0412 | 0.0010 |
| 29 | YGL211W | NCS6 | 42.6040 | 0.0019 |
| 30 | YDL102W | cdc2-7 | 42.5950 | 0.0246 |
| 31 | YMR197C | vti1-11 | 42.5419 | 0.0075 |
| 32 | YGL148W | ARO2 | 42.2665 | 0.0006 |
| 33 | YLR262C | YPT6 | 42.1825 | 0.0057 |
| 34 | YKL110C | KTI12 | 41.6725 | 0.0009 |
| 35 | YLR026C | sed5-1 | 41.6098 | 0.0473 |
| 36 | YDR315C | IPK1 | 41.4007 | 0.0058 |
| 37 | YGL136C | MRM2 | 40.9146 | 0.0025 |
| 38 | YGL130W | ceg1-34 | 40.3700 | 0.0009 |
| 39 | YDR369C | XRS2 | 40.1903 | 0.0417 |
| 40 | YLL006W | MMM1 | 40.0481 | 0.0028 |
| 41 | YNL215W | IES2 | 39.8396 | 0.0264 |
| 42 | YNL141W | AAH1 | 39.2696 | 0.0063 |
| 43 | YDR208W | mss4-103 | 39.0948 | 0.0101 |
| 44 | YMR296C | lcb1-4 | 38.7882 | 0.0299 |
| 45 | YML107C | PML39 | 38.1511 | 0.0012 |
| 46 | YBR171W | SEC66 | 37.9920 | 0.0006 |
| 47 | YGR157W | CHO2 | 37.6673 | 0.0091 |
| 48 | YCR032W | BPH1 | 36.6601 | 0.0122 |
| 49 | YFL039C | act1-129 | 35.2918 | 0.0205 |
| 50 | YOR174W | med4-6 | 35.2918 | 0.0206 |
| 51 | YER013W | prp22-1 | 34.4501 | 0.0042 |
| 52 | YLR249W | yef3-F650S | 34.0535 | 0.0399 |
| 53 | YOR221C | MCT1 | 33.7759 | 0.0003 |
| 54 | YER019W | ISC1 | 33.7416 | 0.0008 |
| 55 | YLL004W | orc3-70 | 33.7003 | 0.0280 |
| 56 | YJR057W | cdc8-2 | 32.8249 | 0.0249 |
| 57 | YDL081C | RPP1A | 32.5448 | 0.0039 |
| 58 | YLR410W | VIP1 | 32.1537 | 0.0043 |
| 59 | YBR237W | prp5-1 | 31.9004 | 0.0019 |
| 60 | YDR062W | lcb2-19 | 31.6883 | 0.0321 |
| 61 | YLL050C | cof1-8 | 31.4206 | 0.0056 |
| 62 | YFL039C | act1-112 | 31.0345 | 0.0371 |
| 63 | YBR095C | RXT2 | 30.9731 | 0.0419 |
| 64 | YLR117C | clf1-1 | 29.8685 | 0.0387 |
| 65 | YOR293W | RPS10A | 29.6764 | 0.0110 |
| 66 | YMR143W | RPS16A | 29.4622 | 0.0025 |
| 67 | YDR087C | rrp1-1 | 28.8367 | 0.0271 |
| 68 | YGL145W | tip20-5 | 28.5626 | 0.0059 |
| 69 | YJR032W | CPR7 | 28.2847 | 0.0030 |
| 70 | YML049C | rse1-1 | 28.1768 | 0.0051 |
| 71 | YLR208W | sec13-1 | 27.8738 | 0.0038 |
| 72 | YDR062W | lcb2-2 | 27.6909 | 0.0128 |
| 73 | YNL302C | RPS19B | 27.6881 | 0.0132 |
| 74 | YDL190C | UFD2 | 26.9511 | 0.0231 |
| 75 | YNR026C | sec12-1 | 26.6261 | 0.0051 |
| 76 | YNL207W | rio2-1 | 25.9212 | 0.0366 |
| 77 | YOR196C | LIP5 | 25.1279 | 0.0382 |
| 78 | YCR081W | SRB8 | 24.9605 | 0.0155 |
| 79 | YMR139W | RIM11 | 24.2426 | 0.0010 |
| 80 | YDR502C | SAM2 | 23.9535 | 0.0056 |
| 81 | YDR145W | taf12-L464A | 23.1961 | 0.0162 |
| 82 | YKR007W | MEH1 | 22.2943 | 0.0468 |
| 83 | YMR123W | PKR1 | 22.1681 | 0.0191 |
| 84 | YBR177C | EHT1 | 20.2462 | 0.0308 |
