## Supplementary file 2 for "Maturation and detoxification of synphilin-1 inclusion bodies regulated by sphingolipids"

**Table S2** Distribution of GO terms among 84 candidate hits with significantly increased Class 3 aggregates via ClueGO analysis

| ID | Term | Ontology Source | P Value | Corrected P Value | Associated Genes (%) | Enrichment | Associated Genes |
| --- | --- | --- | --- | --- | --- | --- | --- |
| GO:0032447 | protein urmylation | Biological Process | 7.44E-05 | 6.10E-03 | 42.86 | 29.78 | NCS6, UBA4, URE2 |
| GO:0098796 | membrane protein complex | Cellular Component | 6.77E-04 | 4.80E-02 | 4.12 | 2.86 | CDC48, EMC1, EMC4, MEH1, MMM1, SEC13, SEC66, SED5, SRP101, VTI1, YME1 |
| GO:0042175 | nuclear outer membrane-endoplasmic reticulum membrane network | Cellular Component | 1.37E-05 | 1.16E-03 | 4.22 | 2.93 | CDC48, CHO2, EMC1, EMC4, IRA2, LCB1, LCB2, MMM1, PKR1, SEC12, SEC13, SEC66, SRP101, TIP20, TSC3, VMA21, VTI1 |
| GO:0005789 | endoplasmic reticulum membrane | Cellular Component | 8.84E-06 | 7.60E-04 | 4.36 | 3.03 | CDC48, CHO2, EMC1, EMC4, IRA2, LCB1, LCB2, MMM1, PKR1, SEC12, SEC13, SEC66, SRP101, TIP20, TSC3, VMA21, VTI1 |
| GO:0044432 | endoplasmic reticulum part | Cellular Component | 2.01E-05 | 1.69E-03 | 4.10 | 2.85 | CDC48, CHO2, EMC1, EMC4, IRA2, LCB1, LCB2, MMM1, PKR1, SEC12, SEC13, SEC66, SRP101, TIP20, TSC3, VMA21, VTI1 |
| GO:0040008 | regulation of growth | Biological Process | 1.77E-04 | 1.36E-02 | 9.38 | 6.51 | LST8, RXT2, TOR2, TSC11, UBA4, VIP1 |
| GO:0001558 | regulation of cell growth | Biological Process | 9.60E-05 | 7.68E-03 | 13.89 | 9.65 | LST8, TOR2, TSC11, UBA4, VIP1 |
| GO:0038201 | TOR complex | Cellular Component | 5.74E-04 | 4.13E-02 | 23.08 | 16.04 | LST8, TOR2, TSC11 |
| GO:0031932 | TORC2 complex | Cellular Component | 2.48E-04 | 1.88E-02 | 30.00 | 20.85 | LST8, TOR2, TSC11 |
| GO:0030950 | establishment or maintenance of actin cytoskeleton polarity | Biological Process | 5.74E-04 | 4.13E-02 | 23.08 | 16.04 | LST8, TOR2, TSC11 |
| GO:0031929 | TOR signaling | Biological Process | 1.42E-04 | 1.11E-02 | 12.82 | 8.91 | LST8, MEH1, SEC13, TOR2, TSC11 |
| GO:1902533 | positive regulation of intracellular signal transduction | Biological Process | 5.18E-04 | 3.78E-02 | 13.79 | 9.58 | MEH1, SEC13, TOR2, TSC11 |
| GO:0002178 | palmitoyltransferase complex | Cellular Component | 1.18E-04 | 9.31E-03 | 37.50 | 26.06 | LCB1, LCB2, TSC3 |
| GO:0008610 | lipid biosynthetic process | Biological Process | 3.76E-04 | 2.78E-02 | 4.78 | 3.32 | CHO2, EHT1, INO4, ISC1, LCB1, LCB2, LIP5, MCT1, TSC11, TSC3 |
| GO:0006665 | sphingolipid metabolic process | Biological Process | 3.49E-04 | 2.62E-02 | 10.64 | 7.39 | ISC1, LCB1, LCB2, TSC11, TSC3 |
| GO:0030148 | sphingolipid biosynthetic process | Biological Process | 8.35E-05 | 6.76E-03 | 14.29 | 9.93 | ISC1, LCB1, LCB2, TSC11, TSC3 |
| GO:0031211 | endoplasmic reticulum palmitoyltransferase complex | Cellular Component | 1.18E-04 | 9.31E-03 | 37.50 | 26.06 | LCB1, LCB2, TSC3 |
| GO:0017059 | serine C-palmitoyltransferase complex | Cellular Component | 4.29E-05 | 3.56E-03 | 50.00 | 34.74 | LCB1, LCB2, TSC3 |
| GO:0035339 | SPOTS complex | Cellular Component | 4.29E-05 | 3.56E-03 | 50.00 | 34.74 | LCB1, LCB2, TSC3 |
