## Supplementary file 3 for "Maturation and detoxification of synphilin-1 inclusion bodies regulated by sphingolipids"

**Table S3** Plasmids used in the present study

| Plasmids | Description | Source |
| --- | --- | --- |
| pYX212-Dsred -SY1 | *2μ, URA3, Amp^R^, TPIp-Dsred-SY1* | Büttner et al, 2010 |
| pYX212-Dsred | *2μ, URA3, Amp^R^, TPIp-Dsred* | Büttner et al, 2010 |
| pYX212-Dsred -SY1-His | *2μ, HIS3, Amp^R^, TPIp-Dsred-SY1* | This study |
| pLCB1-MoBY | *CEN, KanMX^R^, Cm^R^, URA3, LCB1** | Ho et al, 2009 |
| pLCB2-MoBY | *CEN, KanMX^R^, Cm^R^, URA3, LCB2** | Ho et al, 2009 |
| pTSC3-MoBY | *CEN, KanMX^R^, Cm^R^, URA3, TSC3** | Ho et al, 2009 |
| p5472 | *CEN, Cm^R^, URA3* | Ho et al, 2009 |
| pcDNA3.1-GFP-SY1 | pcDNA3.1 (-) vector*, CMVp-GFP-SY1-BGHt* | This study |
| pcDNA3.1-SY1-Flag | pcDNA3.1 (-) vector, *CMVp-SY1-Flag-BGHt* | This study |
| px459-SPTLC2-gRNA1 | px459-CRISPR/Cas9-Puro vector, *U6p-SPTLC2-gRNA1-polIIIt* | This study |
| px459-SPTLC2-gRNA2 | px459-CRISPR/Cas9-Puro vector, *U6p-SPTLC2-gRNA2-polIIIt* | This study |

*Gene expression is controlled by its native promoter and terminator.
