## Supplementary file 4 for "Maturation and detoxification of synphilin-1 inclusion bodies regulated by sphingolipids"

**Table S4** Strains used in the present study

| Strain Name | Genotype and markers | Background/Source |
| --- | --- | --- |
| BY4741 (WT) | *MATa his3Δ1 leu2Δ0 ura3Δ0 met15Δ0* | Dharmacon Inc |
| Y7092+Dsred -SY1 | *MATα* *can1Δ::STE2pr-Sp_his5 lyp1Δ ura3Δ0 leu2Δ0 his3Δ1 met15Δ0 LYS2*+ pYX212-Dsred -SY1 (URA+) | This study |
| Y7092+Dsred | *MATα* *can1Δ::STE2pr-Sp_his5 lyp1Δ ura3Δ0 leu2Δ0 his3Δ1 met15Δ0 LYS2*+ pYX212-Dsred (URA+) | This study |
| BY4741+Dsred -SY1 | BY4741 with pYX212-Dsred -SY1 (URA+) | This study |
| BY4741+Dsred | BY4741 with pYX212-Dsred (URA+) | This study |
| *xxxΔ+*Dsred -SY1 | *MATa xxxΔ::kanMX4 his3Δ1 leu2Δ0 ura3Δ0 met15Δ0+ pYX212-Dsred -SY1* (URA+) | This study |
| *xxxΔ+*Dsred -SY1+p5472 | *MATa xxxΔ::kanMX4 his3Δ1 leu2Δ0 ura3Δ0 met15Δ0 + pYX212-Dsred -SY1-His + p5472* (URA+, HIS3+) | This study |
| *xxxΔ+*Dsred -SY1+ pMoBY | *MATa xxxΔ::kanMX4 his3Δ1 leu2Δ0 ura3Δ0 met15Δ0+ pYX212-Dsred -SY1-His + pMoBY* (URA+, HIS3+) | This study |
| XXX-GFP | BY4741 XXX-GFP (HIS+) | Yeast GFP Clone Collection |
| XXX-GFP+ Dsred -SY1 | BY4741 XXX-GFP + pYX212-Dsred-SY1 (URA+, HIS+) | This study |
